# Schwann cell demyelination is triggered by a transient mitochondrial calcium release through Voltage Dependent Anion Channel 1

**DOI:** 10.1101/581157

**Authors:** Nicolas Tricaud, Benoit Gautier, Gerben Van Hameren, Jade Berthelot, Sergio Gonzalez, Roman Chrast

## Abstract

The maintenance of the myelin sheath by Schwann cells around peripheral nerve axons is essential for the rapid propagation of action potentials. A large number of peripheral neuropathies results for the loss of this myelin sheath, a process called demyelination. Demyelination is a program of cell dedifferentiation characterized by reprograming and several catabolic and anabolic events. This process was best characterized in Wallerian demyelination that occurs following nerve injury. In this model, the earliest well characterized steps are MAPK pathways activation and cJun phosphorylation and nuclear localization starting around 4hrs after nerve injury. Here we show, using *in vivo* imaging of virally-delivered fluorescent probes to mitochondria, that Schwann cell mitochondria pH, motility and calcium are altered as soon as 1hr after nerve injury. Mitochondrial calcium release through VDAC1 mitochondrial channel and mPTP directly induced Schwann cell demyelination via MAPK and c-Jun activation. Decreasing mitochondrial calcium release through VDAC1 silencing or TRO19622 blocking prevented MAPK and cJun activation and demyelination. VDAC1 opening with Methyl Jasmonate induced these cellular mechanisms in absence of nerve injury. Taken together, these data indicate that mitochondria calcium homeostasis through VDAC1 is instrumental in the Schwann cell demyelination process and therefore provide a molecular basis for an anti-demyelinating drug approach.

## Introduction

Schwann cells (SC) are responsible for myelin production in the peripheral nervous system. These cells wrap axons with many turns of compacted and modified plasma membrane in order to form a compact electrically-isolating sheath that allows the fast salutatory propagation of action potentials. Moreover, Schwann cells are thought to be essential for providing neurotrophic and metabolic support to their axons (*1, 2*).

The peripheral myelin is highly plastic. Indeed, in case of peripheral nerve injury, the distal part of the axons degenerate, SC terminate their myelin sheath and enter a dedifferentiation program called Wallerian demyelination (*3*). The myelin is then digested by the SC themselves or by macrophages that are recruited (*4*). When axons have grown back, a re-myelination process then starts to restore a quasi-perfect myelin sheath (*5*). Beside nerve injury, Schwann cell demyelination can also occur due to hereditary and acquired demyelinating diseases of the peripheral nervous system (PNS), which are numerous and affect an increasing number of people (*6*). Acquired demyelinating diseases are not uncommon as they include diabetic peripheral neuropathy (*7*), drug-related peripheral neuropathies, leprosy and peripheral neuropathies of inflammatory etiology (*8*). Hereditary demyelinating diseases of the PNS are rare but remain among the most common hereditary diseases (*9*). While demyelinating peripheral neuropathies are rarely lethal, they range from life threatening to severely affecting life and therefore put a high burden on public health systems. The etiologies of all these acquired and hereditary peripheral diseases are diverse but they all result in demyelination. Thus, an important challenge is to understand the cellular and molecular events that underlie the transition from myelination to demyelination.

Wallerian demyelination represents a relevant model for the study of the SC demyelination program and how it is triggered. After nerve injury one of the first events that occurs in myelinating SC is the activation of MAPK pathways ERK, JNK and P38 (*3*). Then shortly after cJun is phosphorylated and accumulates in the nucleus (*10*). However, what triggers these events remains unclear. Recent studies show that mitochondria dysfunctions are involved in an increasing number of demyelinating neurodegenerative diseases (*11, 12*). Myelin sheath mitochondria have been shown to be essential for neuron homeostasis but their morphological and physiological properties remain elusive. It is well known that patients suffering from mitochondrial multisystem disorders often show a peripheral neuropathy in addition to other more debilitating symptoms (*13*). Myelinating Schwann cells (SC) of patients suffering these diseases display features of demyelination as well as abnormal mitochondria. More directly, some acquired demyelinating PNS diseases appear to be linked to defects in mitochondrial functions. Indeed a side effect of some anti-HIV drugs is demyelinating peripheral neuropathy (*14*). In addition, this drug-induced pathology and the diabetic peripheral neuropathy have been linked with perturbation of mitochondrial functions and to mitochondrial stress and defects in PNS (*15*). Taken together, these studies suggest that mitochondria can play an essential role in peripheral demyelinating diseases.

The goal of this project was to investigate the role of mSC mitochondria during Wallerian demyelination. Using *in vivo* fluorescent probes and multiphoton imaging, we show here that mSC mitochondria release their calcium in the cytoplasm through VDAC1 and the mPTP and this leads to the activation of the demyelination program.

## Results

### Live imaging of mSC mitochondria and fluorescent probes validation

The right sciatic nerves of 2 months old mice were injected with a solution of adenovirus or AAV expressing different fluorescent probes that infected myelinating Schwann cells *in vivo* (*16–18*). Three weeks later, mice were anesthetized, their sciatic nerve exposed and placed under a multiphoton microscope allowing mSC mitochondria time-lapse imaging in physiological conditions (**Fig. 1A**) (*16*). We used mito-Dsred2 (*16*) to measure mitochondrial size and movements, mito-Sypher (*19*) for mitochondrial matrix pH, mito-GCaMP2 (*20*) for mitochondrial matrix calcium content and the same probe without the peptide targeting to mitochondria to assess of cytoplasmic calcium content (cyto-GCaMP2). Mito-Dsred2 probe was previously validated *in vivo* (*16*). Mito-GCaMP2 and cyto-GCaMP2 were validated by incubating infected nerve in successive baths of calcium chelator EDTA and calcium donor CaCl2 in a mild saponin detergent (**Fig. 1B and C**). Mito-Sypher was validated by incubating infected nerves in successive bathes of sodium azide and ammonium chloride (**Fig. 1D**), which are respectively inhibitor and activator of the probe activity (*21*). In all cases, probes were found to be functional in myelinating SC *in vivo* and no demyelination occurred.

**Figure 1.**
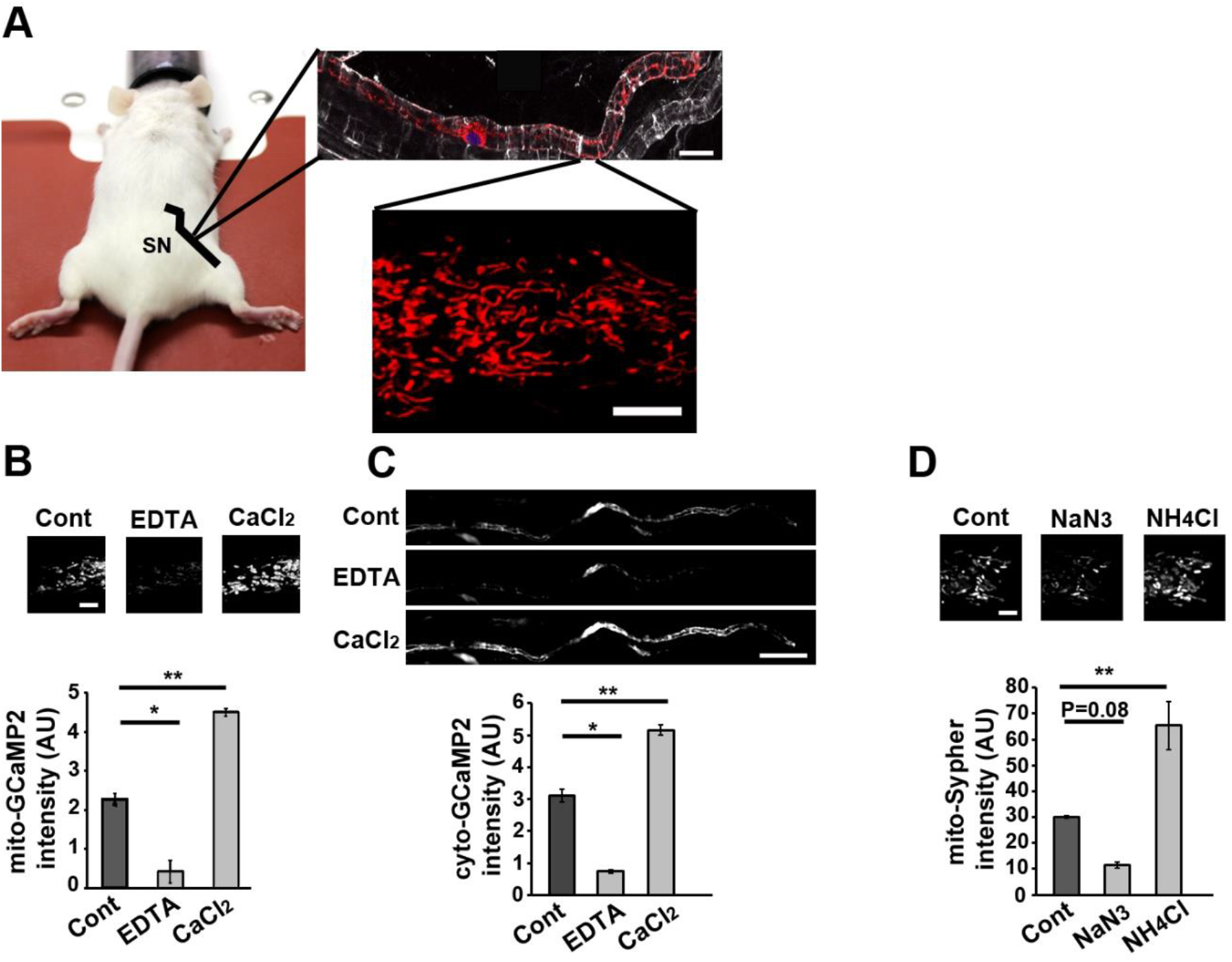
Live imaging technique and fluorescent probes validation. **A**- Schematic representation of the imaging technique used in this work. The sciatic nerve (SN) of anesthetized mice illustrated by the black line was exposed under the lens of a multiphoton microscope. Myelinating SC, stained for E-cadherin (white), contains mitochondria labelled through adenovirus-delivered mito-Dsred2 expression (red) in particular around the cell nucleus labelled with TOPRO3 (blue)(scale bar= 10μm). This technique allows the selective *in vivo* multiphoton imaging of mitochondria in some part of living SC (scale bar=5μm). All mice were 8 to 11 weeks old. **B**- Mouse sciatic nerves expressing mito-GCaMP2 were incubated in a bath containing Leibovitz’s L15 medium without (Cont) or with EDTA (1mM) or Calcium chloride (CaCl_2_, 100 μM). Probe fluorescence was imaged (upper panels pictures, scale bar=5μm) and measured (lower panel graph). **C**- Mouse sciatic nerves expressing cyto-GCaMP2 were incubated in a bath containing Leibovitz’s L15 medium without (Cont) or with EDTA (1mM) or Calcium chloride (CaCl_2_, 100 μM). Probe fluorescence was imaged (upper panels pictures, scale bar=50μm) and measured (lower panel graph). **D**-Mouse sciatic nerves expressing mito-Sypher were incubated in a bath containing Leibovitz’s L15 medium without (Cont) or with sodium azide (Na3, 3mM pH 3.2) or ammonium chloride (NH4Cl, 30 mM pH 8). Probe fluorescence was imaged (upper panels pictures, scale bar=5μm) and measured (lower panel graph). Error bars show SEM. Statistical tests are one-way ANOVA comparing Control with each other condition. n=2 experiments for B and C and 3 experiments for D.

### Sciatic nerve injury induces mitochondrial calcium release in the cytoplasm, mitochondria slowdown and matrix alkalization in mSC

Next, sciatic nerves of anesthetized animals were placed under the microscope and the probe fluorescence was recorded for around 20 minutes to setup a stable baseline of fluorescence. Then, nerves were injured to induce Wallerian demyelination through carefully crushing around 5 mm upstream of the imaging area. Changes occurring in probe-labelled myelinating SC mitochondria downstream of the crush were recorded over 5 hrs. Mito-Dsred2 probe reported a slow but significant decrease of the mitochondrial speed starting 120 minutes after the injury (**Fig. 2A**). Looking at the mitochondrial length, we classified them as short (0.1 to 1μm), medium (1 to 3μm) and long (>3μm)(**Fig. 2B**). We did not observe any significant change in the frequency of each size category (**Fig. 2B**), suggesting mitochondria slowed down after the trigger of demyelination without fragmenting as seen previously after animal death (*16*). Mito-Sypher reported a significant and persistent increase in pH starting 120 minutes after the injury (**Fig. 2C**), showing an alkalization of the mitochondrial matrix. Mito-GCaMP2 fluorescence showed a sharp decrease of mitochondrial calcium around one hour after the injury (**Fig. 3A**). This loss of calcium in mitochondria timely correlated with a surge of calcium in the myelinating SC cytoplasm reported by cyto-GCaMP2 (**Fig. 3B**), indicating mitochondrial calcium was released in the cytoplasm. This was just a pulsed release of calcium as mitochondrial and cytosolic calcium levels quickly returned to their basal initial levels. However, mitochondrial calcium significantly increased 2 hours after the injury indicating a hypercalcemia of the mitochondrial matrix. Taken together, mitochondrial calcium release to the cytoplasm occurred before the increase in matrix pH and the slowdown of mitochondria movements (**Fig. S1**). Matrix hypercalciema then occurred following this change in pH (**Fig. S1**).

**Figure 2.**
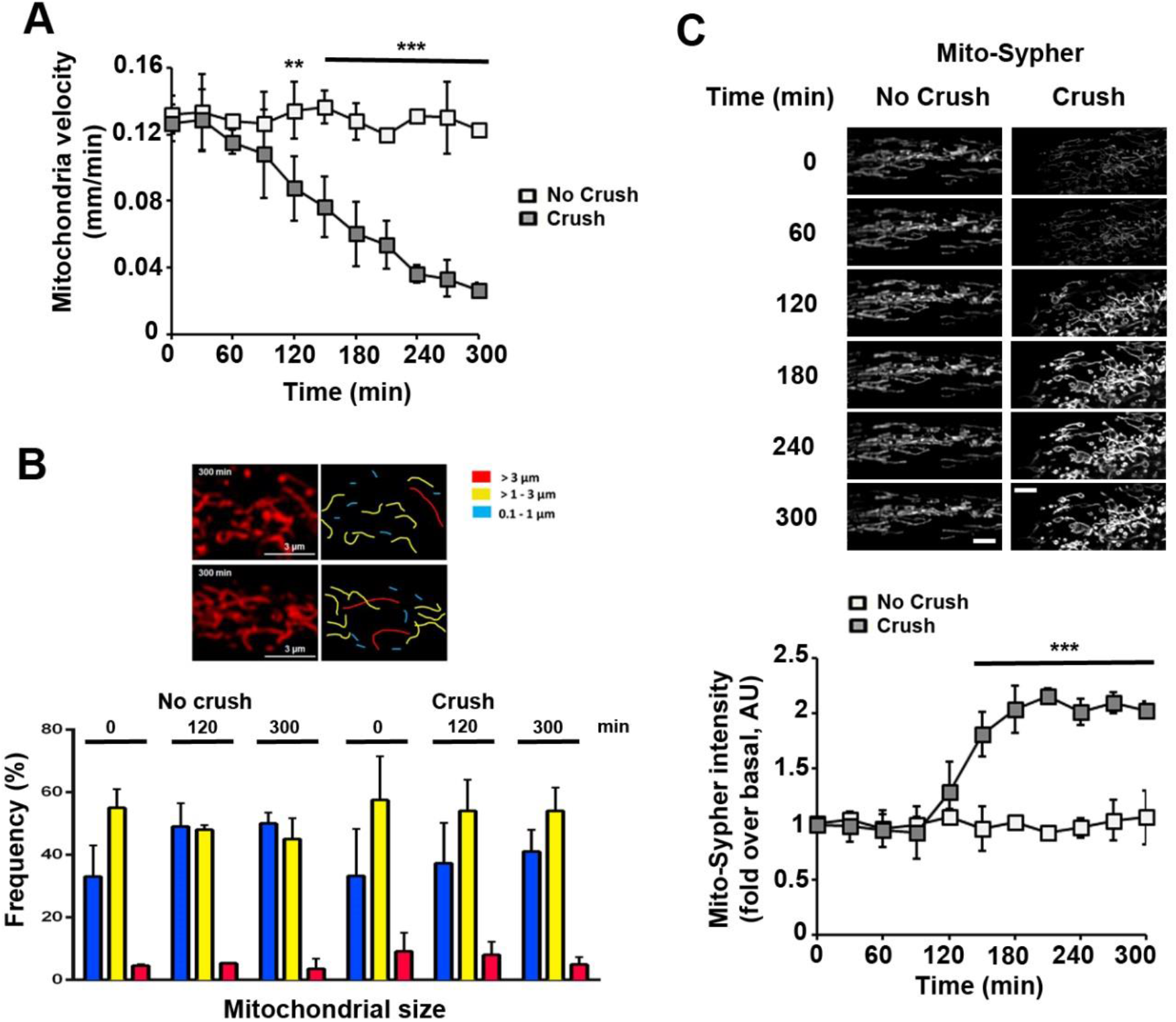
Mitochondrial mobility and pH change during Schwann cell demyelination. **A**- Mitochondrial velocity in myelinating SC of control (No crush, n=2 mice) and crushed (Crush, n=5 mice) nerves was measured using adenovirus-delivered mito-DsRed2. Velocity is shown in mm traveled in one minute. **B- Upper panels**: Representative *in vivo* images of SC mitochondria (left panels) showing how mitochondrial length is characterized and quantified (right panels) in non-crushed and crushed nerves. **Lower panel**: Frequency histogram of mitochondrial size in control (No crush, n=2 mice) and crushed (Crush, n=5 mice) conditions at 3 successive time points. **C- Upper panels**: Representative *in vivo* images of SC mitochondria labeled with adenovirus-delivered mito-Sypher probe in non-crushed (No Crush) and crushed nerves (Crush) at successive time points. Scale bar= 5μm. Lower panel: the average fluorescence intensity of the probe was measured for more than 100 mitochondria at the successive indicated time points in non-crushed (No Crush, n=3 mice) and crushed nerves (Crush, n=4 mice). The probe fluorescence intensity is normalized over the basal condition before crush. Error bars indicate SEM. Statistical tests are repeated measures two-way ANOVA Sidak *post hoc* test comparing non-crushed to crushed nerves values. All mice were 8 to 11 weeks old.

**Figure 3.**
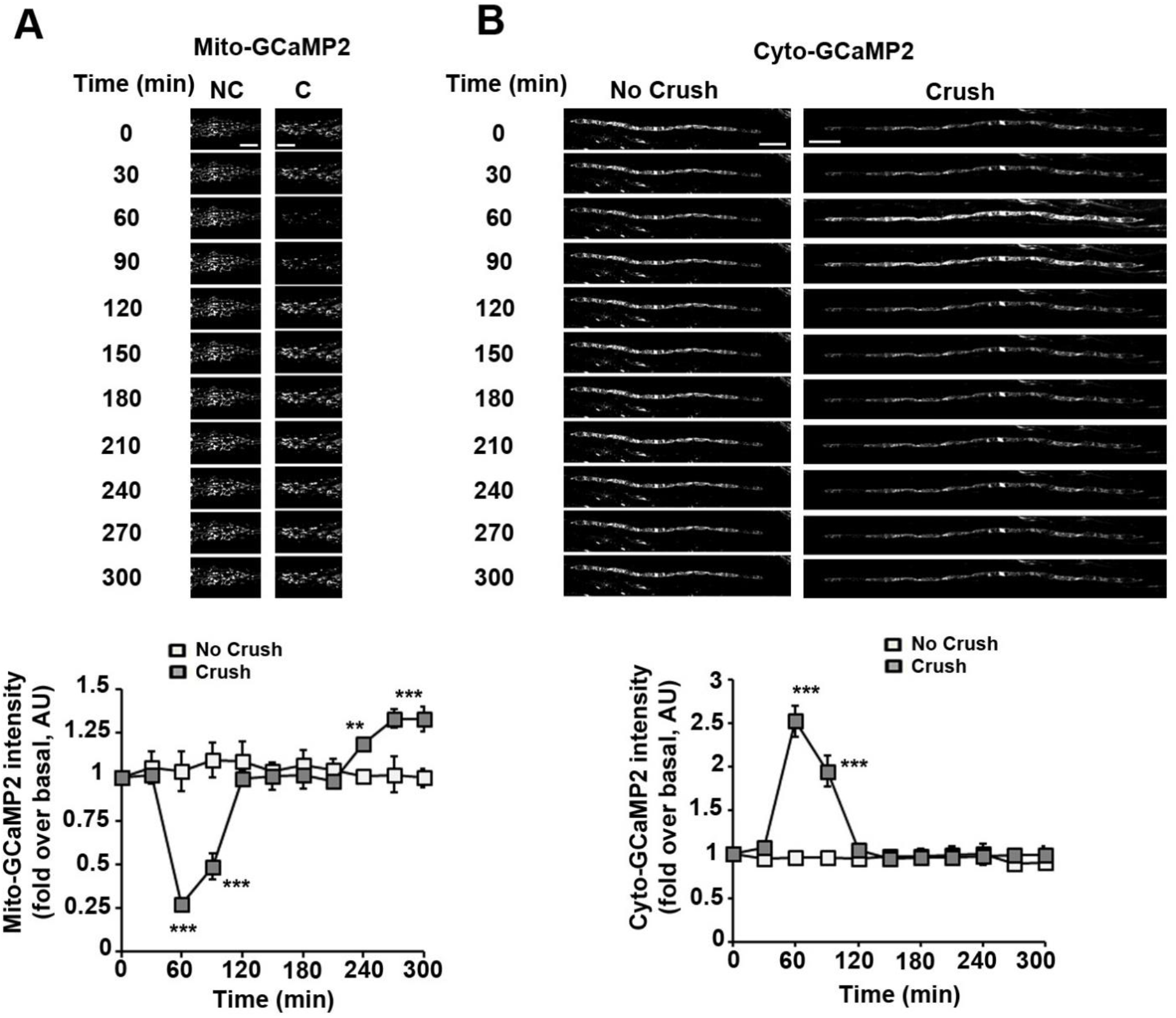
Mitochondrial and cytoplasmic calcium dynamics in SC following nerve injury. **A- Upper panels**: Representative *in vivo* images of SC mitochondria labeled with adenovirus-delivered mito-GCaMP2 probe in non-crushed (No Crush) and crushed nerves (Crush) at successive time points. Scale bar= 5μm. **Lower panel**: the average fluorescence intensity of the probe was measured for more than 100 mitochondria at the successive indicated time points in non-crushed (No Crush, n=3 mice) and crushed nerves (Crush, n=5 mice). The probe fluorescence intensity is normalized over the basal condition before crush. **B- Upper panels**: Representative *in vivo* images of SC mitochondria labeled with adenovirus-delivered cyto-GCaMP2 probe in non-crushed (No Crush) and crushed nerves (Crush) at successive time points. Scale bar= 50μm. **Lower panel**: the average fluorescence intensity of the probe was measured for more than 100 mitochondria at the successive indicated time points in non-crushed (No Crush, n=3 mice) and crushed nerves (Crush, n=4 mice). The probe fluorescence intensity is normalized over the basal condition before crush. Error bars indicate SEM. Statistical tests are two-way ANOVA Sidak *post hoc* test comparing non-crushed to crushed nerves values. All mice were 8 to 11 weeks old.

### VDAC1 and mPTP are responsible for mitochondrial calcium release in the cytoplasm

Mitochondrial calcium dynamics changed very quickly both in mitochondria and cytoplasm after triggering Wallerian demyelination. As mitochondrial calcium is known to act a signaling event in the cell, we next investigated the molecular mechanisms that mediate the mitochondrial calcium release. VDAC1, Voltage-Dependent Anion-selective Channel 1, is a porin ion channel (*22*) highly expressed in mSC outer mitochondrial membrane (*23*) and regulating mitochondrial calcium release, cell signaling, apoptosis and dedifferentiation (*24, 25*). We characterized two shRNAs selectively silencing mouse VDAC1 expression in Schwann cell line MSC80 (**Fig. 4A**) and in myelinating SC *in vivo* (**Fig. 4B**). A control shRNA with no target in mammalian cells was also characterized. Viral vectors expressing both the fluorescent probes and the shRNAs were used to transduce myelinating SC *in vivo*. We observed that both shRNAs targeting VDAC1 significantly reduced mitochondrial calcium decrease and the concomitant cytoplasmic calcium increase after nerve injury while the control shRNA had no effect (**Fig. 4C**). This indicated that VDAC1 is required for the efflux of calcium from the matrix to the cytoplasm in these conditions. In order to confirm, we also tested TRO19622 (Olesoxime), a neuroprotective drug that has been shown to bind VDAC(*26, 27*), as inhibitor of calcium release through VDAC1. TRO19622 inhibited both mitochondrial calcium decrease and cytoplasmic calcium increase in myelinating SC after injury (**Fig. 4D**). Hexokinase binding is the main endogenous inhibitor of VDAC in the cell (ref) and the release of hexokinase from VDAC through methyl jasmonate (MJ) induces several mitochondrial and cellular modifications (*28*). When nerves expressing mito-GCaMP2 were treated for two hours with MJ, without any injury, mitochondrial calcium significantly decreased (**Fig. 4E**). This showed that VDAC opening was sufficient to release of calcium even in absence of injury.

**Figure 4.**
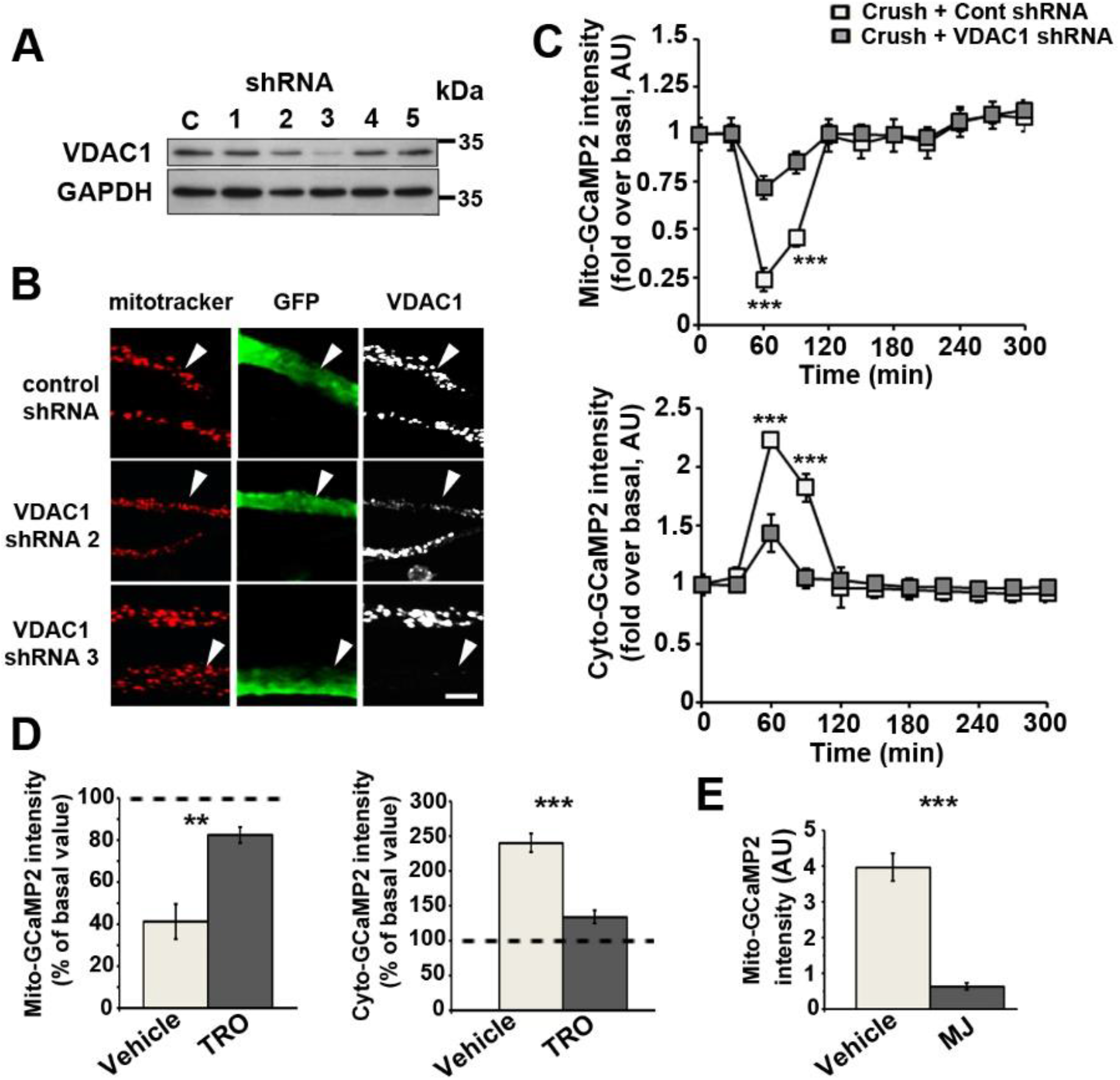
VDAC1 mediates mitochondrial calcium release during SC demyelination. **A**-MSC80 SC were transfected with plasmids expressing puromycin and a control shRNA with no mammalian target or shRNAs specifically targeting mouse VDAC1 under a U6 promoter. After a selection with puromycin, cells were lysed and their VDAC protein expressions were analyzed by WB. GAPDH is used as loading control. **B**- AAV virus expressing VDAC1 shRNAs 2 or 3 in addition to GFP were injected in the sciatic nerve of mice. Three weeks later animals were sacrificed and the expression of VDAC in infected SC was analyzed by immunostaining on teased fibers. Infected cells expressing control shRNA and GFP (green, arrowheads) show mitochondria (mitotracker, red) harboring VDAC (white) when cells expressing VDAC1 shRNAs show a reduced expressing of VDAC. Scale bar=5 μm. **C**- AAV virus expressing VDAC1 shRNAs 2 or 3 in addition to mito- or cyto-GCaMP2o were injected in the sciatic nerve of mice. The average fluorescence intensity of the mito-GCaMP2 (upper panel) and cyto-GCaMP2 (lower panel) probes was measured for more than 100 mitochondria at the successive indicated time points in non-crushed (No Crush, n=3 mice) and crushed nerves (Crush, n=4-5 mice). The probe fluorescence intensity is normalized over the basal condition before crush. Error bars indicate SEM. Statistical tests are two-way ANOVA Sidak *post hoc* test comparing crushed with Control shRNA to crushed with VDAC1 shRNAs conditions. All mice were 8 to 11 weeks old. **D**- Sciatic nerves transduced with viruses expressing mito-GCaMP2 (left panel, n=3 for both TRO and vehicle) or cyto-GCaMP2 (right panel, n=3 for both TRO and vehicle) were injected with 2μl of 20μM TRO19622 (TRO) solution or vehicle 30 minutes before nerve injury and then live imaging. Probes fluorescence intensities were measured before injury and then 60 minutes after injury at the peak of mitochondrial calcium release. Values at t=60 min are presented in percentage of basal values before crush. The dotted lines indicate 100% of the basal values. Error bars indicate SEM. Statistical tests are two-tailed Student t-test. All mice were 8 to 11 weeks old. **E**- Sciatic nerves transduced with viruses expressing mito-GCaMP2 (left panel, n=89 and 109 mitochondria for vehicle and MJ respectively, in 4 mice each) were injected with 1μl of 57μM methyl jasmonate (MJ) solution or vehicle 2 hrs before live imaging. Probes fluorescence intensities were measured at that time on regions of interest with a similar number of mitochondria to limit variability. Error bars indicate SEM. Statistical tests are two-tailed Student t-test.

However, as VDAC is located in the outer mitochondrial membrane, the involvement of another channel crossing the inner mitochondrial membrane is required to explain the release of matrix calcium in the cytoplasm. VDAC participates to the formation of the mitochondrial Permeability Transition Pore, mPTP (*29*), which crosses the inner mitochondrial membrane. To check the involvement of mPTP in the mitochondrial calcium release following nerve injury, we selectively blocked mPTP activity using cyclosporine A (*30*). The concentration required to block mPTP was determined using a saturating concentration of auranofin, a drug that opens mPTP (*31*), combined with an increasing concentration of cyclosporine A (**Fig. S2**). With this concentration, cyclosporine A was able to prevent the release of mitochondrial calcium following nerve injury (**Fig. 5**), showing that mPTP is indeed involved in this process in addition to VDAC.

**Figure 5.**
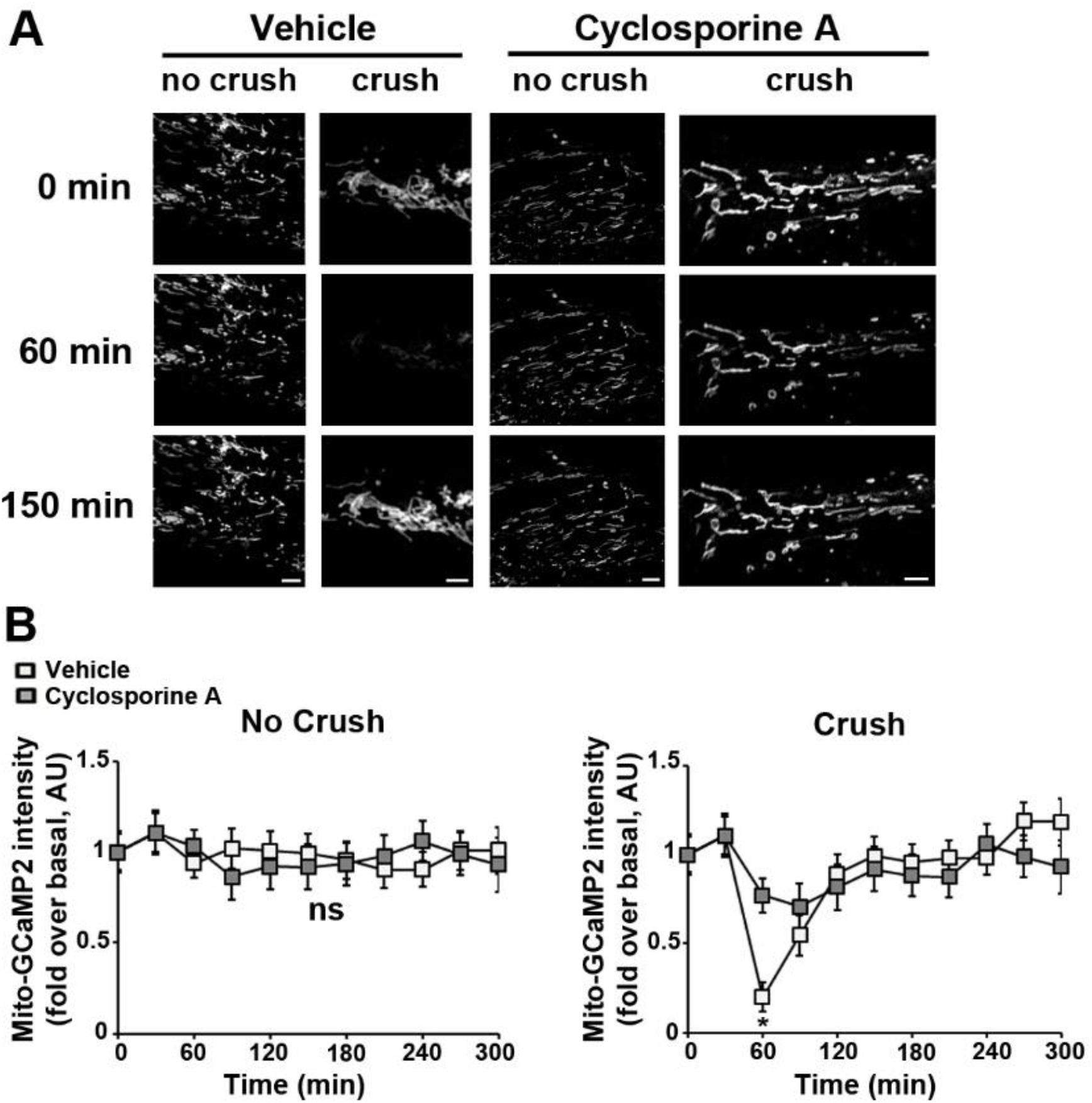
Blocking mPTP with cyclosporine A prevent mitochondrial calcium release in SC following nerve injury. **A**- Representative *in vivo* images of SC mitochondria labeled with adenovirus-delivered mito-GCaMP2 probe in non-crushed (No Crush) and crushed (Crush) nerves at successive time points following treatment with 500 μm cyclosporine A (right panels) or vehicle (left panels). Scale bar= 3μm. **B- Left panel**: Average fluorescence intensity of mito-GCaMP2 was measured for more than 100 mitochondria at the successive indicated time points in non-crushed nerves treated with cyclosporine A or vehicle. **Right panel**: Average fluorescence intensity of mito-GCaMP2 was measured for more than 100 mitochondria at the successive indicated time points in crushed nerves treated with cyclosporine A or vehicle. The probe fluorescence intensities for each graph are normalized over the basal condition before crush or at the first imaging time point for non-crushed nerves. Error bars indicate SEM. Statistical tests are two-way ANOVA Sidak’s *post hoc* test comparing cyclosporine A treated and vehicle treated conditions. n=3 animals for each condition. All mice were 8 to 11 weeks old.

### The release of mitochondrial calcium via VDAC1 induces Schwann cell demyelination *in vivo* via MAPK and cJun activation

As nerve crush initiates SC demyelination, we investigated if the release of mitochondrial calcium could activate pathways known to be involved in the demyelination process: ERK1/2, p38 and JNK (*32–34*) and cJun phosphorylation and nuclear enrichment (*35*). Phospho-ERK1/2, phospho-p38, phospho-JNK and phospho-cJun significantly increased 4 and 12 hours after sciatic nerve injury and blocking VDAC through TRO19622 treatment strongly reduced these pathways activation (**Fig. 6A and S3**). Conversely, MJ treatment increased demyelination pathways activation in non-crushed nerve (**Fig. 6A and S3**). Additionally, we observed a strong reduction of nuclear localization of phospho-cJun when VDAC1 was silenced (**Fig. 6B**) or blocked though TRO19622 (**Fig. 6C**). The opening of VDAC through MJ treatment induced phospho-cJun nuclear enrichment in absence of injury (**Fig. 6D**). Taken together this shows that VDAC opening induces the activation of the demyelination pathways in sciatic nerves and the enrichment of phospho-cJun in the nucleus of SC.

**Figure 6.**
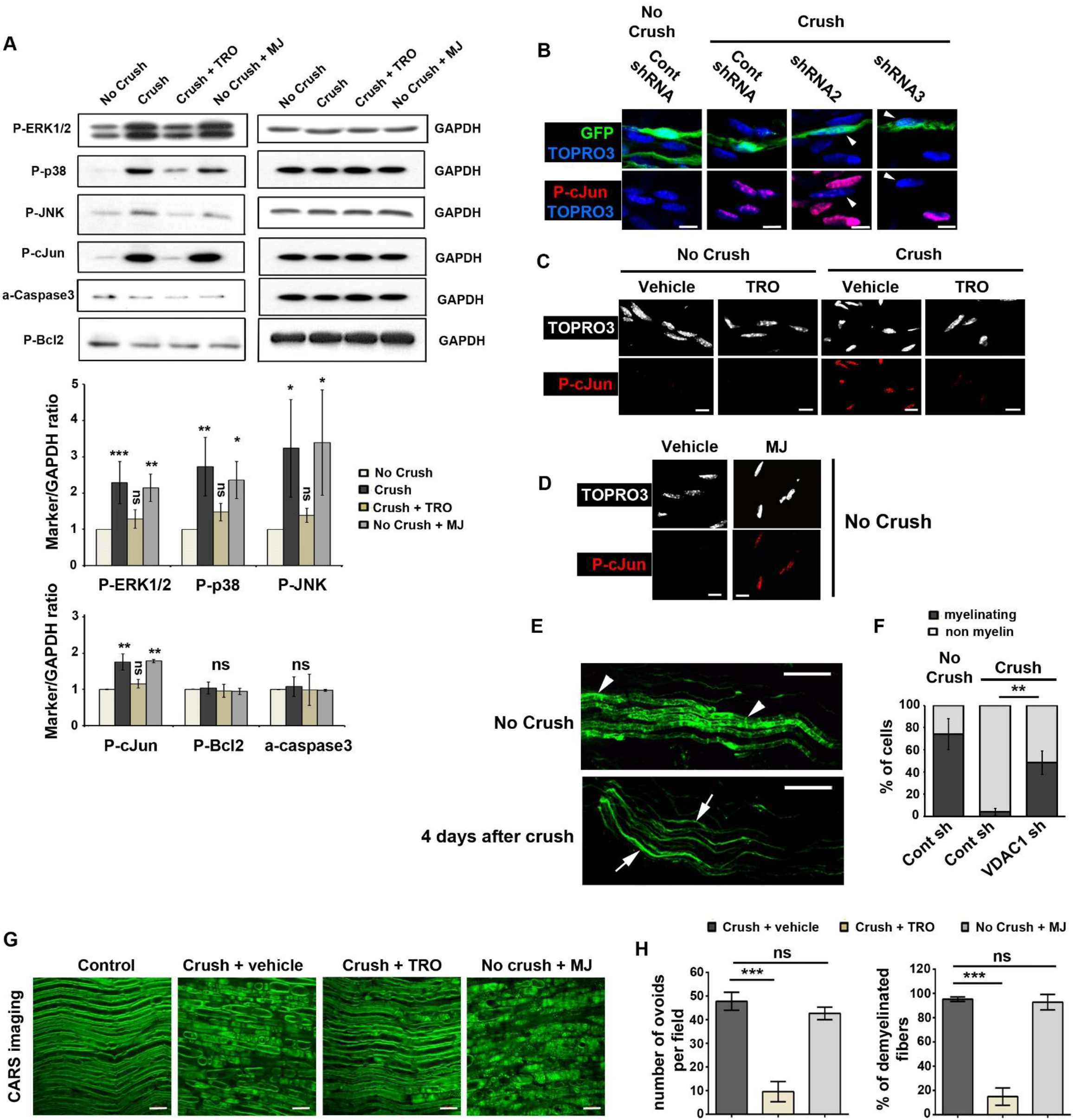
Demyelination is decreased after treatment with TRO19622 and spontaneously induced after treatment with MJ. **A- Upper panels**: Western blot analysis for phospho-ERK1/2 (P-ERK1/2), phospho-p38 (P-p38), phospho-JNK (P-JNK), phospho-cJun (P-cJun), activated cleaved-caspase3 (a-caspase3) and phospho-Bcl2 (P-Bcl2) in sciatic nerve of mice without injury (No crush), 4 hrs after injury (Crush), 4 hrs after injury with TRO19622 treatment 30 min before injury (Crush + TRO) or without injury but 4 hrs after methyl jasmonate treatment (No crush + MJ). GAPDH was used as loading control. **Lower panels**: WB results analyzed by densitometry and normalized on GAPDH respective values on the same blot. As similar changes were observed at 4 and 12hrs after injury (see Fig. S3 for 12hrs representative images), data from both experiments were pooled except for P-cJun, for which changes were consistent but much higher at 12hrs. So only 4hrs data are presented. All data are presented as fold over No crush condition. Error bars show SD. Statistical tests are one-way ANOVA followed by a Dunnett’s multiple comparison *post hoc* tests for each marker. n = 2 (P-cJun) to 4 mice. ns= non-significant. **B**- Immunohistochemistry for phospho-cJun (P-cJun, red) and nuclear TOPRO3 (blue) on teased fibers of mouse nerve transduced with virus expressing control shRNA or shRNA 2 or 3 targeting VDAC1 in addition to GFP (green). Without injury (No crush) non infected cells or cells expressing control shRNA and GFP show no P-cJun in their nucleus. Four days after injury (Crush), non-infected cells and cells expressing control shRNA and GFP express P-cJun in their nucleus while cells expressing shRNA2 or 3 and GFP show low amount of P-cJun in their nucleus (arrowheads). Scale bars=10μm. **C**- Immunohistochemistry for phospho-cJun (P-cJun, red) and nuclear TOPRO3 (white) on teased fibers of mouse nerve without injury (No crush) or 4 days after injury (Crush). In both cases, animals were treated with TRO19622 (TRO) or vehicle, once intraperitoneally 10 hrs before injury, once intrasciatically 30 min before injury and then intraperitoneally for 4 consecutive days. Scale bars=10μm. **D**- Immunohistochemistry for phospho-cJun (P-cJun, red) and nuclear TOPRO3 (white) on teased fibers of mouse nerve without injury (No crush) 4 days after treatment with methyl jasmonate (MJ) or vehicle. Scale bars=10μm. **E**- Representative images of the morphological features occurring in myelinating SC during demyelination 4 days after nerve injury. These cells express virally-delivered GFP (Green). Arrowheads show myelinating SC and arrows non-myelin forming cells. Scale bars=100μm. **F**- Quantification of myelinating and non-myelinating SC frequency in nerves transduced with AAV-expressing Control shRNA (Cont sh) or shRNAs 2 or 3 targeting VDAC1 (VDAC1 sh) in addition to GFP with (Crush) or without nerve injury (No crush). Nerves were collected 4 days after the nerve injury and GFP-positive cells were counted as myelinating or non-myelin forming as shown in Fig. 6E. Data are represented as mean ±SD. Statistical significances were determined using a two-tailed Student’s t-test. n= 5 (No Crush, Cont sh), 4 (Crush, Cont sh) and 4 (Crush, VDAC1 sh) mice. **G**- Representative CARS images of sciatic nerves collected and fixed by a 4% PFA solution in non-injured control animals (Control), in animals 4 days after crush nerve injury and after vehicle (Crush + vehicle) or TRO19622 (Crush + TRO) treatment or in animals 4 days after injection of MJ in the sciatic nerve (No Crush + MJ). Scale bars= 20μM. **H**- The number of ovoid per field (left) and the percentage of demyelinated fibers per field (right) were measured on the CARS images. Data are presented as mean ±SEM. n=5 (Crush + vehicle), 5 (Crush + TRO) and 4 (No Crush + MJ) animals. Statistical analysis shows one-way ANOVA followed by a Tukey’s *post hoc* test.

To go further, we investigated the demyelination using typical defects induced in the cellular morphology of mSC as described previously (*36*). Myelinating SC have homogeneously large and long morphology *in vivo* and, after demyelination, these cells acquire a non-myelin forming cells features characterized by a heterogeneous thin and split morphology with fine extensions (**Fig 6E**). Following the labelling of myelinating SC with a virus expressing GFP and control or VDAC1 shRNAs, the right sciatic nerve of anesthetized mice were exposed and crushed. Seven days later the injured nerves were collected and fixed. After teasing of the fibers, GFP-expressing cells were analyzed using confocal microscopy. Nerve injury induced demyelination features in a large majority of cells expressing GFP and control shRNA while, when cells expressed GFP and VDAC1 shRNA, significantly fewer cells showed a demyelination phenotype (**Fig. 6F**), showing that VDAC1 silencing protects cells from demyelination.

Finally, we tested the anti-demyelinating effect of TRO19622 and the pro-demyelinating effect of MJ *in vivo* using CARS imaging of myelin as read-out for demyelination. Coherent Anti-stokes Raman Scattering (CARS) is a non-linear microscopy approach that allows imaging lipids, hence myelin, *in vivo* or *ex vivo* (*37, 38*). First, two months old mice were treated with 3mg/kg of TRO19622 or vehicle subcutaneously for 5 consecutive days. The second day these mice were anesthetized, their right sciatic nerves exposed and crushed. Four days later, the injured nerves were collected, fixed and analyzed using CARS imaging. In crushed nerves treated with vehicle, the myelin sheath was fragmented in ovoids compared to non-injured nerves (**Fig. 6G**) due to the collapse of the Schmidt-Lantermann incisures, which represents the first morphological feature of demyelination (*3, 4*). This fragmentation was quantified or by counting the number of ovoids or by counting the percentage of demyelinating fibers (**Fig. 6H**). However, when animals were treated with TRO19622, this fragmentation was absent or very preliminary (**Fig. 6G, quantified in 6H**) showing that inhibiting mitochondrial calcium-release prevents demyelination *in vivo*. Moreover, when animals were injected intrasciatically with MJ a spontaneous demyelination occurred in absence of crush (**Fig. 6G and H**) confirming that VDAC opening is sufficient for the initiation of demyelination.

## Discussion

While demyelinating peripheral nerve diseases include a large spectrum of acquired and inherited disabling diseases, mechanisms of SC demyelination remain elusive. Peripheral nerve demyelination does not result from cell death but from myelinating SC dedifferentiation (*39*). So demyelination is a cellular program in which the myelinating SC enters upon specific signals such as axonal injury in a trauma. These triggering signals are transduced in the cell and drive the activation of MAPK demyelination pathways (p-ERK1/2, p-P38 and p-JNK activation) followed by the recruitment of phosphorylated cJun in the nucleus. Here we describe an earlier step, the release of mitochondrial calcium and changes in mitochondria physiology.

Using a viral approach to express fluorescent probes in mitochondria of myelinating SC in the sciatic nerve of living mice and a multiphoton microscope for time-lapse live imaging, we show that the release of mitochondrial calcium occurs as soon as one hour after inducing demyelination by nerve injury. This release is followed, in this specific order, by a burst of calcium in the cytoplasm, the slowing of mitochondrial movements, the increase of mitochondrial pH and mitochondrial hypercalcemia (**Fig. S1**). Using silencing and activation/inhibition with drugs, we show that VDAC1 is responsible for the release of mitochondrial calcium in the cytoplasm. This pulse of mitochondrial calcium through VDAC1 activates the known demyelination pathways ERK1/2, p38 and JNK leading to cJun phosphorylation in the nucleus, which characterizes the demyelination program in myelinating SC (*35*). However several other unclear cellular processes are also engaged and this will result in the collapse of the cell structure, the breakdown of the myelin (*3, 4*) and the recruitment of macrophages to help to clear myelin debris (*4*). Once demyelination is completed, around 5 days after crush of sciatic nerve, then dedifferentiated SC will be able to remyelinate axons.

The release of calcium by mitochondria is the earliest step recorded after nerve injury and our data indicate that this step is necessary and sufficient for triggering the demyelination program. How this burst of mitochondrial calcium through VDAC1 channels activates ERK1/2, p38 and JNK pathways is not clear. However mitochondrial calcium release is an essential cell signaling triggering death and differentiation (*40*). Moreover cytoplasmic calcium stimulates ERK1/2 (*41*) and JNK (*42*) activity. Several studies also reported an association of ERK1/2, p38 and JNK and their direct activators such as Raf, Sab, and MKK4 with mitochondria (*43*). So it is likely that mitochondrial calcium release in the cytoplasm directly activates MAPK demyelination pathways and these cascades pathways propagate the demyelination signal all over the cell. Nevertheless other mechanisms cannot be excluded such as a production of reactive oxygen species (ROS) in calcium-activated mitochondria to stimulate ERK1/2, p38 and JNK pathways (*43, 44*). Whatever is the precise mechanism this mitochondrial signaling after nerve injury never reached the cell death level as cell-death related pathways such as caspase 3 and Bcl2 were not activated. Mitochondrial signaling is therefore able to generate a differentiation process instead of cell death, as previously reported in myoblasts (*45*).

VDAC has numerous binding partners that controls its permeability and in particular hexokinase (HK). HK binding to VDAC reduces the permeability of the pore notably to calcium (*24*). In SC methyl jasmonate, a compound that uncouples HK from VDAC (*28*), induced mitochondrial calcium release and demyelination showing that HK is essential to control demyelination. Intriguingly, mutations in HK gene are responsible for demyelinating CMT4G disease (*46*), suggesting that an increased permeability of VDAC to calcium is the cause of the disease. On the opposite we show that TRO19622, a compound that binds to VDAC (*27*), blocks mitochondrial calcium release through VDAC, therefore preventing demyelination.

Mitochondrial calcium release through VDAC has other consequences that may also contribute to the demyelination program. First the slowdown of mitochondrial motility we observed after nerve crush is likely to be due to the increase of cytoplasmic calcium around mitochondria. Indeed, calcium alters the activity of molecular motors that mediate mitochondrial movements in the cell (*47*). Second, mitochondrial pH is linked to cytoplasmic calcium concentration (*48*). So, when mitochondria release calcium their pH changes. However, while hypocalcemia is transitory, the pH changes on the long term (at least for 5 hours) suggesting that mitochondrial respiratory activity is increased (*49*). As ATP controls the uptake of calcium from endoplasmic reticulum (ER) (*50*), the mitochondrial hypercalcemia occurring 4 hours after nerve crush reflects also an increase in respiratory activity. Taken together this suggests that mitochondrial calcium release after nerve crush profoundly changes mitochondria physiology in mSC, making them more active. How mitochondrial activity and metabolic changes participate to the demyelination program is a very interesting open question.

## MATERIAL AND METHODS

### Animal housing

Immunodeficient strain CB17/SCID mice (Janvier Labs, France) were kept in the animal house facility of the Institute for Neurosciences of Montpellier in ventilated and clear plastic boxes, subjected to standard light cycles (12 h to 90 lux light, 12 h dark). The care, breeding and use of animals followed the animal welfare guidelines of the “Institut National de la Santé et de la Recherche Medicale” (INSERM), under the approval of the French “Ministère de l’Alimentation, de l’Agriculture et de la Pêche”, Approval Number CEEA-LR-11032).

### Cloning

Plasmid dsRed2-mito7 (Clontech #55838) was cut using NheI and NotI enzymes and then treated with DNA polymerase I Large Klenow fragment to isolate mito-dsRed2 cDNA. After purification, it was cloned into pAdtrack-CMV (Quantum Biotechnologies, Inc.) or pAAV-MCS (Cell Biolabs, Inc.) plasmids under the control of a CMV or a CAG promoter respectively. pcDNA3.1 mito-GCaMP2 (kindly provided by Dr. X. Wang, Peking University, China) was cut or with HindIII and EcoRV to be cloned into a pShuttle-CMV (Quantum Biotechnologies, Inc.) or with BamHI and EcoRV to be cloned into a pAAV-MCS vector under the control of a CMV or a CAG promoter respectively. Then, the GCaMP2 probe cDNA (without mito sequence) was cut using HindIII and EcoRV to be cloned into a pAAV-MCS and to pShuttle-CMV vector. The probe mito-SypHer (kindly provided by Dr. J.C. Jonas, Université Catholique de Louvain, Belgium) was cut using NheI and NotI enzymes and treated with DNA polymerase I Large Klenow fragment. After purification, it was cloned into pAAV-MCS under the control of a CAG promoter. The mouse VDAC1-specific shRNA sequence 2 **GTTGGCTATAAGACGGATGAACT** (Sigma-Aldrich, Ref. #TRCN0000012391), the VDAC1 shRNA sequence 3 **ACCAGGTATCAAACTGACGTTCT** (Sigma-Aldrich, Ref. #TRCN0000012392) or the shRNA control (dsRed2) **AGTTCCAGTACGGCTCCAA** or (GFP) **CAAGCTGACCCTGAAGTTC** were first cloned separately into a pSicoR vector (Addgene, Ref. 11579) under the control of a U6 promoter using HpaI and BstEII enzymes then, the U6-VDAC1-shRNA sequences were then cut using ApaI and BstEI to be cloned into a pAAV-CMV-GFP vector (Cell Biolabs, Inc.), the pAAV-mito-GCaMP2, the pAAV-GCaMP2, the pAAV-mito-dsRed2 or the pAAV-mito-SypHer previously described. All clones were validated by sequencing.

### Viral particles production

Adenoviral particles production was described in He et al., 1998 (*51*). Briefly, for adenovirus production pAdtrack vector containing the constructs was recombined with pAdeasy1 vector in the Adeasy1 BJ5183 bacteria strain (Stratagene). The isolated adenoviral DNA was cut with PacI enzyme and transfected in HEK 293 cells using Lipofectamine 2000. The initial production of adenovirus was followed by 3 rounds of amplification. Finally freeze thaw cycles are used to harvest the viral particles from cells, then purified using cesium chloride gradients. To produce high-titer adeno-associated virus (AAV10), three 15 cm dishes of 70 – 80 % confluent HEK293T cells were transfected with 71 μg of pAAV expression vector, 20 μg of pAAV10 capsid and 40 μg of pHelper (Cell Biolabs, Inc.). 48 h later transfection, the medium was collected, pooled and centrifuged 15 min at 2000 rpm to spin down floating cells. In parallel, cells were scraped and collected in PBS. Then, cells were lysed using dry ice/ethanol bath and centrifuged 15 min at 5000 rpm to discard cell debris. The cleared supernatant and the cleared medium were pooled and filtrated using a 0.22 μm filter. The viral solution was filtrated through a cation-exchange membrane Mustang S acrodisc (Pall Corporation) to deplete empty particles and later, filtrated through an anion-exchange membrane Mustang Q acrodisc (Pall corporation) to retain AAV viral particles. Then, viruses were eluted and concentrated using centrifugal concentrators Amicon tube. Usual titer is around 10^11^ PFU/ml. For further details see Okada et al., 2009 (*52*).

### *In vivo* virus injection in the sciatic nerve

Five to seven weeks old mice were anesthetized with isoflurane inhalation and placed under a Stemi2000 microscope (Zeiss). Incision area was shaved and cleaned using betadine solution. After incision, the *gluteus superficialis* and *biceps femoris* muscles were separated to reveal the cavity traversed by the sciatic nerve. The nerve was lift out using spatula and a thin glass needle filled with viral solution (8 μl) was introduced into the nerve with a micromanipulator. This solution was injected over 30 min with short pressure pulses using a Picospritzer III (Parker Hannifin) coupled to a pulse generator. After injection, the nerve was replaced into the cavity, the muscles were readjusted, and the wound was closed using clips (for further details see Gonzalez et al., 2014 (*18*)).

### Sciatic nerve set up under multiphoton microscope

Eight to eleven weeks old injected mice were anesthetized with 5% of isoflurane a 1.5% of oxygen into the anesthesia system box (World Precision Instruments, Ref. EZ-B800) for 5 minutes. Then, mice were placed in anesthesia mask and the incision area was shaved and cleaned with betadine and ethanol 70% solution. Incision was realized using a scalpel and the skin was retracted using forceps in order to expose the *gluteus superficialis* and *bíceps femoris* muscles. Next, the connective tissue that connects both muscles was cut and the sciatic nerve was gently lifted out using a spatula. A flexible bridge was slide below sciatic nerve and it was placed into a plastic first pool fixed to the bridge. Mice were placed under the multiphoton microscope, the bridge was fixed using magnetic brackets to avoid physiological movement and mouse legs were fixed using clippers. Microscope dark box temperature is controlled to 37 °C during all time-lapse imaging. Finally, a second pool was fixed to the first pool using a drop of agarose low melting 3% (Promega, Ref. V2111) in Leibovitz’s L15 medium (Gibco Life Technologies) and filled with deionized water to immerse the objectives 20x or 63x (Carl Zeiss Microscopy). Sciatic nerves were crushed using serrated forceps at least 5mm above the imaging area. Five successive crushes were performed at different angles to maximize the demyelination all around the nerve.

### Multiphoton image acquisition and analysis

All time-lapse images were obtained with a multiphoton microscope Zeiss LSM 7 MP OPO. Mitochondria motility images were acquired by time-lapse recording of one image every five minutes during five hours using mito-Dsred 2 labelled mitochondria (excitation light wavelength 920 nm). GCaMP2 and SypHer probe images were acquired by time-lapse recording of one image every fifteen minutes during five hours (excitation light wavelength 985 nm). Images were acquired with constant laser intensity (1%), 100 ms of acquisition time, 512×512 resolution and 10 images per z-stack (20 μm). Images were saved in Zeiss .czi format and processed using Image J program. Analysis were done as described in Gonzalez et al., 2015(*16*).

### Immunohistochemistry

The right sciatic nerves of 8 weeks old mice were crushed as described previously. Four to twelve hours later these nerves were dissected and washed in L15 medium, fixed in Zamboni’s fixative for 10 min at room temperature, washed in PBS, and incubated in successive glycerol baths (15, 45, 60, 66% in PBS) for 18 to 24 h each, before freezing at −20 °C. The nerves were cut in small pieces in 66% glycerol and the perineurium sheet removed. Small bundles of fibers were teased in double-distilled water on Superfrost slides, dried overnight at room temperature, and the slides stored at −20 °C. For immunostaining, the teased fibers were incubated for 1 h at room temperature in blocking solution (10% goat serum, 0.2% TritonX100, and 0.01% sodium azide in PBS). Then, the samples were incubated with anti-ECCD2 primary mouse antibody (1/100, BD biosciences, Ref. 610181), anti-phosphoS63-c-jun primary mouse antibody (1/200, BD Biosciences, Ref. 558036), anti-c-jun primary mouse antibody (1/200, BD Biosciences, Ref. 610326), anti-VDAC primary rabbit antibody (1/100, Cell Signaling, Ref. 4866) or/and MitoTracker Red (1/1000, Molecular Probes, Ref. M7515) in blocking solution overnight at 4 °C. The next day, the samples were washed in PBS and incubated for 1 h at room temperature with secondary donkey antibodies coupled to Alexa488, Alexa 594 or Alexa647 (1/1600, Molecular probes) and TOPRO3 iodide (50 μM, Invitroge, Ref. T3605). Finally the samples were washed in PBS and mounted in Immu-mount (Thermo Scientific). Images were acquired at room temperature using a 20x or 40x objective, a Zeiss confocal microscope LSM710, and its associated software.

### Intranerve drug administration

TRO19622 drug (Tocris bioscience, Ref. 2906) was diluted in ethanol to 20mM then, diluted in sterile PBS to 20μM. Intrasciatic TRO19622 treatment for live imaging experiments was realized by intra sciatic nerve injection of 2 μl of the 20μM drug solution using a Hamilton syringe or a glass needle held on a micromanipulator as described for the virus injection. This injection occurred 30 minutes before multiphoton imaging or nerve injury followed by multiphoton imaging. Methyl jasmonate treatment was realized by injection of 7μl of MJ commercial solution (30 μmol) into the sciatic nerve using a glass needle held on a micromanipulator two hours before multiphoton image acquisition or 4 days before CARS imaging. Auranofin (Sigma Aldrich, Ref. A6733) was diluted in DMSO and 4 pmol (2 μl) were injected in the sciatic nerve using a Hamilton syringe during multiphoton image acquisition. Cyclosporine A (Sigma Aldrich, Ref. 30024-25MG) was diluted to 100 mM in ethanol then, diluted in sterile PBS to 500 μM. Cyclosporine A treatment was realized by intra sciatic nerve injection of 4 μl of solution (200 pmol) using a glass needle and micromanipulator 30 minutes before multiphoton image acquisitions. Ethanol for TRO19622 or PBS for methyl jasmonate and Cyclosporine A were used as vehicle.

### Systemic drug administration

TRO19622 (Bio-techne, Ref.2906) was diluted to 0,3mg/ml in Cremophore EL/dimethylsulfoxide/ethanol/phosphate buffer saline (CDEP, 5/5/10/80 v/v). TRO19622 treatment (3mg/kg) was performed daily by subcutaneous injection. Control mice were injected the same way with CDEP (5/5/10/80, v/v).

### CARS imaging and Image Analysis

Four days after MJ injection or nerve crush, animal were sacrificed and their sciatic nerves were collected, washed by PBS and fixed for 1h in 4% PFA solution at room temperature. All CARS images were obtained as described previously (*37*) with a two-photon microscope LSM 7 MP coupled to an OPO (Zeiss, France) and complemented by a delay line (*53*). A ×20 water immersion lens (W Plan Apochromat DIC VIS-IR) was used for image acquisition. At least 3 wide-field images were acquired per nerve and per condition. For each image, the number of myelin ovoids and the percentage of degenerated fibers were counted using ZEN software (Zeiss).

### Cell culture and transfection

Mouse Schwann cells MSC80 (ExPASy, CVCL_S187) were grown in DMEM (Dulbecco’s modified Eagle’s medium) (Gibco Life Technologies) supplemented with 2 mM L-glutamine, 100 U/ml^−1^ penicillin/streptomycin and 5% (v/v) heat-inactivated fetal bovine serum (all supplements were from Invitrogen). Cells were maintained at 37°C in an atmosphere of 5% CO_2_, and were passaged when they were 80–90% confluent, twice a week. Five 15 cm dishes of 70 – 80 % confluent mouse Schwann cells were separately transfected with 30 μg of pLKO.1-puro vector containing five commercial VDAC1 shRNA (Sigma-Aldrich, sh1#TRCN0000012388, sh2#TRCN0000012389, sh3#TRCN0000012390, sh4#TRCN0000012391 and sh5#TRCN0000012392) using 80 μl of Lipofectamine 2000 (Invitrogen) and 1,5 ml of Opti-Mem (Gibco Life Technologies). After 7 h, the medium was changed to a fresh complete culture medium enriched with 2mM of glutamine. 48h after transfection cells were treated with 0.5 μg/ml of Puromycin (Gibco Life Technologies) for selection. Antibiotic was maintained in cell medium during one week. Then, cells were collected for western blot.

### Protein extraction and Western blotting

Transfected cells were washed in PBS, lysed in lysis buffer (10 mM Tris, pH 7.4, 150 mM NaCl, 1% Triton X-100, 0.1% SDS, 0.5% sodium-deoxycholate, 1mM EDTA, 50mM NaF, 1mM NaVO4, protease inhibitor cocktail (Sigma-Aldrich)) for 15 min on ice, and centrifuged at 14,000 rpm at 4°C to pellet cell debris. Sciatic nerves were dissected from 8 weeks old mice with or without crush, washed in PBS and directly fixed with 4% of PFA for 10 minutes. After removal of the epineurium and perineurium, the nerves were homogenized by sonication in lysis buffer. Cellular debris was removed by centrifugation at 13,000 g for 5 min at 4ºC and protein was quantified by the bicinchoninic acid method using bovine serum albumin as a standard. Then, samples were denatured at 98 °C, loaded on 10% SDS-PAGE, and transferred on PVDF membranes for immunoblotting. Antibody against the phospho-S63-cJun (1/100, BD Biosciences, Ref. 558036) was from mouse and antibodies against VDAC (1/1000, Cell Signaling, Ref. 4866), phospho-Thr183/Tyr185-SAPK/JNK (1/1000, Cell Signaling, Ref. 9251), phospho-Thr202/Tyr204-p44/42 MAPK (ERK 1/2) (1/1000, Cell Signaling, Ref. 9101), phospho-Thr180/Tyr182-p38 (1/1000, Cell Signaling, Ref. 9211), Cleaved Caspase-3 (1/1000, Cell Signaling, Ref. 9661), total non-phosphorylated JNK (1/1000, Cell Signaling, Ref. 9252) were from rabbit. Antibody against phospho-S87-bcl2 was from goat (1/500, Santa Cruz Biotechnology, Ref. sc-16323).

### Validation of fluorescent probes and anesthesia control

Mouse sciatic nerves expressing mito-GCaMP2 or mito-SypHer were isolated three weeks after AAV particles infection. Nerves were washed in PBS and incubated in Leibovitz’s L15 medium (Gibco Life Technologies) for 3 hours at 37°C in an atmosphere of 5% CO_2_. Sciatic nerve infected with mito-SypHer probe was treated separately with sodium azide 3mM solution at pH 3.2 (Sigma-Aldrich, Ref. S2002) or with ammonium chloride 30 mM solution at pH 8 (Sigma-Aldrich, Ref. A4514) for 5 minutes. Sciatic nerve infected with mito-GCaMP2 was treated separately with calcium chelator EDTA 1mM solution (Sigma-Aldrich, Ref. ED255) or calcium chloride 100 μM solution (Sigma-Aldrich, Ref. S3014) and saponine 20 μg/μl solution (Sigma-Aldrich, Ref. S4521) for 5 minutes. Mito-SypHer and mito-GCaMP2 probe intensities were quantified at 985 nm using multiphoton microscope.

### Data and statistical analysis

Data are represented as mean ±SEM or ±SD. Statistical significances were determined using a two-tailed Student’s *t* test, one-way ANOVA followed by a Dunnett’s multiple comparison *post hoc* test or by a Tukey’s multiple comparison *post hoc* test and two-way ANOVA followed by a Sidak’s multiple comparison *post hoc* test. Significance was set at * P < 0.05, ** P < 0.01, or *** P < 0.001. ns indicates non-significant differences (p > 0.05). n indicates the number of independent experiments.

## SUPPLEMENTAL INFORMATION

### AUTHOR CONTRIBUTIONS

NT wrote the paper; BG, GvH, SG and JB performed experiments; BG, GvH, SG, RC and NT analyzed data and NT supervised the project.

The authors declare no competing financial interests

## ACKNOWLEDGMENTS

We thank H. Boukhaddaoui (Montpellier RIO Imaging Platform) and the Animal Facility of the INM for help and technical assistance. We would like to thank Dr. J.C. Jonas (Université Catholique de Louvain, Belgium) for mito-sypHer probe, Dr. X. Wang (Peking University, China) for mito-GCaMP2 probe and Dr G. Lenaers for his support. This work was supported by the European Research Council grant (FP7-IDEAS-ERC 311610) and an INSERM - AVENIR grant to NT and The Neuromuscular Research Association Basel, Swedish StratNeuro program, Swedish Research Council grant (#2015-02394) to RC.

