## Supplemental Material for "Schwann cell demyelination is triggered by a transient mitochondrial calcium release through Voltage Dependent Anion Channel 1"

#### FIGURE S1

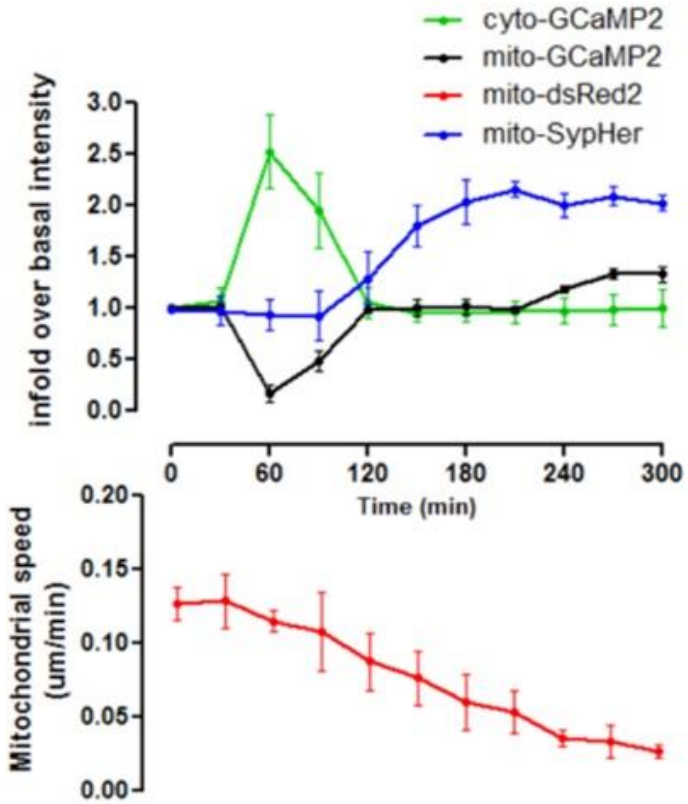


**Figure S1. Succession of events occurring in myelinating SC when Wallerian demyelination is triggered**

We plotted the dynamics of mito-GCaMP2, cyto-GCaMP2, mito-Sypher probes fluorescence and mito-Dsred2 labelled mitochondria velocity on the same time frame. This shows that the first event to occur is the release of calcium from the mitochondria to the cytoplasm (between 30 and 120 min). At the same time mitochondrial velocity starts to drop regularly to reach immobility by around 300 min. Mitochondrial pH increases (>120 min) to reach a plateau by around 180 min. Then mitochondrial calcium increases (>180 min) reaching a plateau by around 270 min.

### FIGURE S2

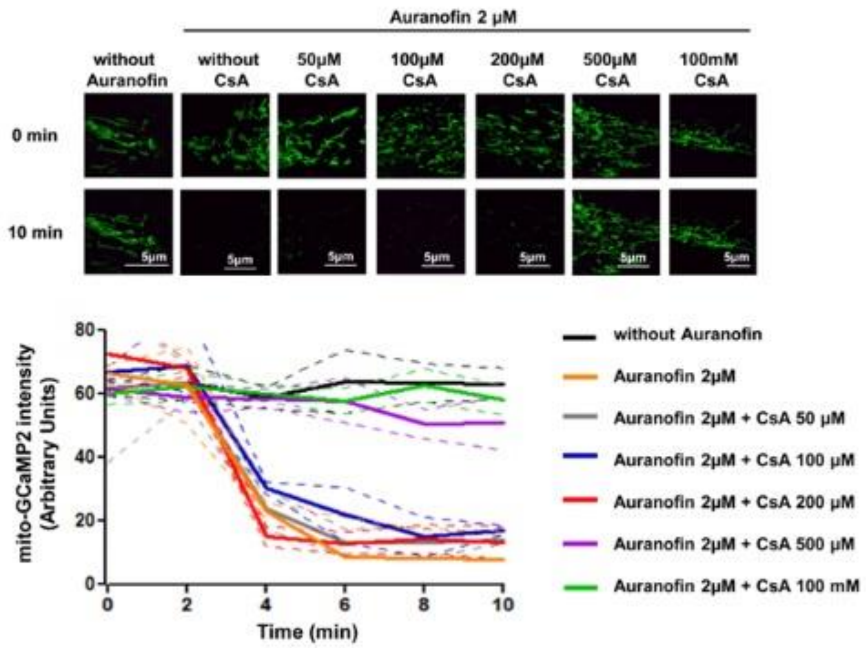


**Figure S2. 500 µM cyclosporine A blocks mPTP opening induced by 2µM auranofin in SC *in vivo***

Mice expressing mito-GCaMP2 in SC were anesthetized and mitochondrial calcium dynamics in SC were analyzed by live two-photon imaging. Cyclosporine A (CsA) was injected in the imaged nerve 30 minutes before starting imaging and then 2 µm auranofin was injected to open mPTP and release mitochondrial calcium. When cyclosporine A concentration is too low (50, 100 and 200µM) auranofin is able to open mPTP and release mitochondrial calcium which decrease the probe fluorescence. Starting at 500µM cyclosporine A is able to block mPTP opening following auranofin injection.

**Upper panel**. Representative images of the mito-GCaMP2 probe fluorescence in the different conditions.

**Lower panel**. The mean mito-GCaMP2 fluorescence intensity was measured for more than 100 mitochondria at different time points for each conditions. n=2 mice.

### FIGURE S3

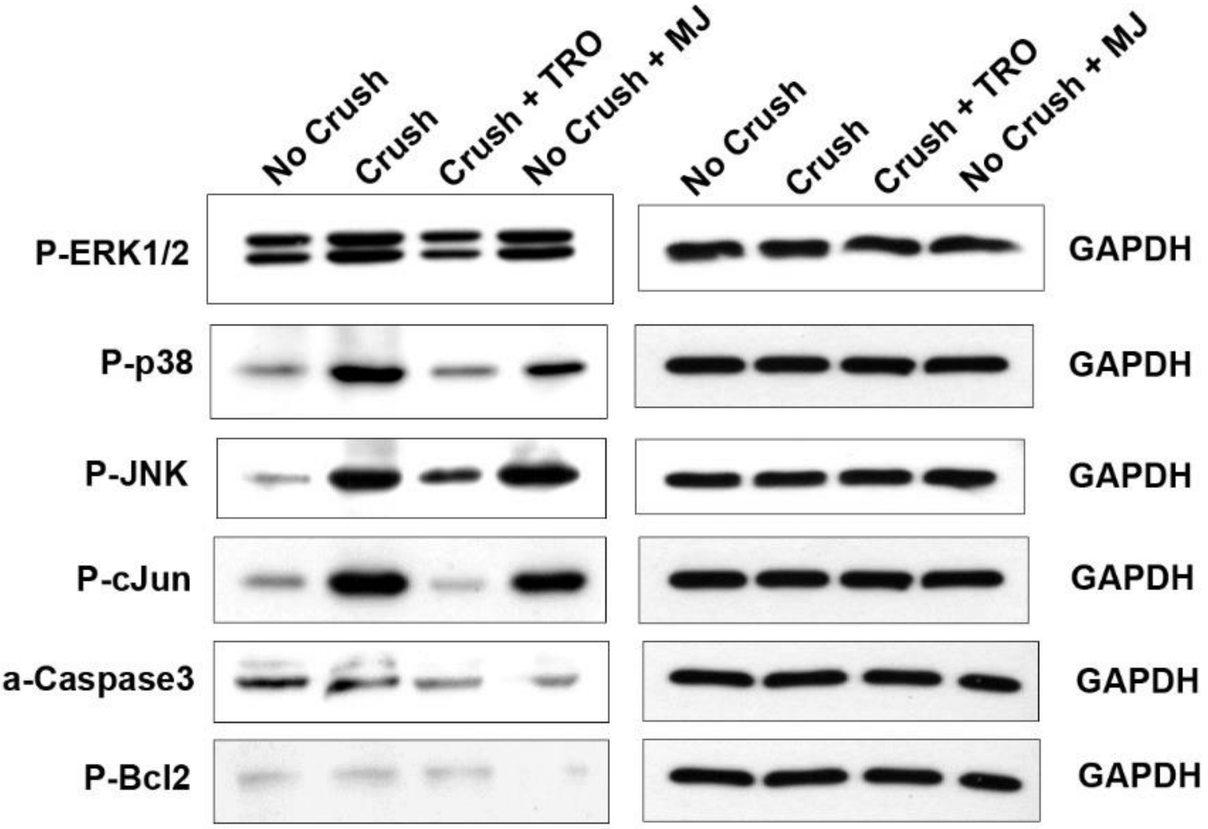


**Figure S3. Activation of the demyelination pathways 12 hrs after injury**

Western blot analysis for phospho-ERK1/2 (P-ERK1/2), phospho-p38 (P-p38), phospho-JNK (P-JNK), phospho-cJun (P-cJun), activated cleaved-caspase3 (a-caspase3)and phospho-Bcl2 (P-Bcl2) in sciatic nerve of mice without injury (No crush), 12 hrs after injury (Crush), 12 hrs after injury with TRO19622 treatment 30 min before injury (Crush + TRO) or without injury but 12 hrs after methyl jasmonate treatment (No crush + MJ). GAPDH was used as loading control.
